# Alterations in large-scale resting-state network nodes following transcranial focused ultrasound of deep brain structures

**DOI:** 10.1101/2024.08.26.609720

**Authors:** Stephanie M. Gorka, Jagan Jimmy, Katherine Koning, K. Luan Phan, Natalie Rotstein, Bianca Hoang-Dang, Sabrina Halavi, Norman Spivak, Martin M Monti, Nicco Reggente, Susan Y. Bookheimer, Taylor Kuhn

**Affiliations:** Department of Psychiatry and Behavioral Health, The Ohio State University Wexner Medical Center, 370 W. 9th Avenue, Columbus, OH 43210; Department of Psychiatry and Biobehavioral Sciences, The University of California, Los Angeles, 760 Westwood Blvd, Los Angeles, CA, 90095; Department of Psychology, The University of California, Los Angeles, 2191 Franz Hall, Los Angeles, CA, 90095; Institute for Advanced Consciousness Studies, 2811 Wilshire Blvd Ste. 590, San Monica, CA, 90403

**Author notes:** **Corresponding Author:** Taylor Kuhn, PhD, Department of Psychiatry and Biobehavioral Sciences, University of California, Los Angeles, Los Angeles, CA, United States, 760 Westwood Blvd, Los Angeles, CA, 90095. **Funding:** UCLA Staglin IMHRO Center for Cognitive Neuroscience Pilot Funding (TK). UCLA Semel Institute for Neuroscience and Behavior (TK).

**Keywords:** transcranial focused ultrasound, resting-state functional connectivity, amygdala, entorhinal cortex

## Abstract

**Background:** Low-intensity transcranial focused ultrasound (tFUS) is a brain stimulation approach that holds immense promise for the treatment of brain-based disorders. Several studies in humans have shown that tFUS can successfully modulate perfusion in focal sonication targets including the amygdala; however, limited research has explored how tFUS impacts the function of large-scale neural networks.

**Objective:** The aim of the current study was to address this gap and examine changes in resting-state connectivity between large-scale network nodes using a randomized, double-blind, within-subject crossover study design.

**Methods:** Healthy adults (n=18) completed two tFUS sessions, 14 days apart. Each session included tFUS of either the right amygdala or the left entorhinal cortex (ErC). The inclusion of two active targets allowed for within-subjects comparisons as a function of the locus of sonication. Resting-state functional magnetic resonance imaging was collected before and after each tFUS session.

**Results:** tFUS altered resting-state functional connectivity (rsFC) within and between rs-network nodes. Specifically, pre-to-post sonication of the right amygdala modulated connectivity within nodes of the salience network (SAN) and between nodes of the SAN and the default-mode network (DMN) and fronto-parietal network (FRP). A decrease in SAN to FPN connectivity was specific to the amygdala target. Pre-to-post sonication of the left ErC was found to modulate connectivity between the dorsal attention network (DAN) and FPN and DMN. An increase in DAN to DMN connectivity was specific to the ErC target.

**Conclusion:** These preliminary findings may suggest that tFUS induces neuroplastic changes beyond the immediate sonication target.

## Introduction

Transcranial focused ultrasound (tFUS) is a novel approach to brain stimulation that holds immense promise for the treatment of brain-based disorders (Pasquinelli et al., 2019; Stern et al., 2021). At high intensities, tFUS can be used to locally ablate brain tissue and provide irreversible treatment (di Biase et al., 2021; Moosa et al., 2019) and at low intensities, tFUS can be used to transiently modulate functioning of specific brain regions by non-invasively applying acoustic energy (Tyler et al., 2006; Dell’Italia et al., 2022). Unlike other neuromodulation approaches, low intensity tFUS can reach deep brain structures with high spatial precision to inhibit or enhance neural activation (Romanella et al., 2020). By manipulating ultrasound parameters, it is also possible to change brain function without causing tissue damage (Spivak et al., 2022). Low intensity tFUS therefore overcomes many of the limitations of existing neuromodulation techniques and is being increasingly explored as a novel treatment approach for a variety of neurological and psychiatric disorders.

tFUS has been applied to a variety of brain regions including somatosensory and visual cortices, insula, thalamus, and striatum (Legon, 2021; Monti et al, 2016; Cain, 2021). Recent studies have also begun to target the amygdala, a brain region known to mediate threat and emotion processing (Ohman, 2005; Folloni et al., 2019; Kuhn et al, 2023; Chou et al., 2024). Three decades of neuroimaging research indicates that hyperactivity of the amygdala is involved in the pathophysiology of internalizing disorders including anxiety and post-traumatic stress disorder (Ressler & Nemeroff, 2000; Gerin et al., 2019). Early studies show that tFUS of the amygdala can modulate amygdala perfusion and alleviate symptoms of anxiety (Zielinski et al., 2021; Mahdavi et al., 2023). For example, a case study of an individual with treatment-resistant generalized anxiety disorder (trGAD) demonstrated that tFUS of the amygdala resulted in immediate decreases in anxiety symptoms (Zielinski et al., 2021). In a larger cohort of patients with trGAD, eight weekly tFUS sessions targeting the amygdala similarly resulted in significant decreases in anxiety symptoms, with 64% of patients achieving clinically significant benefit (Mahdavi et al., 2023). Converging research therefore suggests that tFUS has therapeutic potential and the safety, efficacy, and neural mechanisms of tFUS should be further explored.

How tFUS modulates neural processes is still unclear (Dell’Italia et al., 2022). Studies clearly indicate that tFUS increases or decreases blood oxygen level dependent (BOLD) signal depending upon sonication parameters (Kuhn et al., 2023; Chou et al., 2024). Indeed, a study by our group found that tFUS of the amygdala selectively increased perfusion in the right amygdala but not the left amygdala, without engaging the auditory cortex (Kuhn et al., 2023). How these focal changes influence the rest of the brain is still an area under active investigation. Using BOLD data collected simultaneously with the tFUS sonication experiment, we found decreases in functional connectivity between the right amygdala and posterior cingulate, anterior cingulate, medial prefrontal and posterior parietal regions in a sample of healthy older adults (Kuhn et al., 2023). A separate study by Chou and colleagues (2024) examined changes in resting-state functional connectivity (rsFC) in healthy adults before and after active or sham tFUS of the left amygdala. Chou et al. (2024) reported that active tFUS resulted in decreased amygdala-insula and amygdala-hippocampal rsFC, and increased amygdala-ventromedial prefrontal cortex rsFC.

Existing studies provide important initial evidence that tFUS changes FC between the sonication target and other areas of the brain. However, no study to date has directly examined tFUS-related changes to neural networks beyond the target. The brain is organized into multiple distributed (large-scale) systems including the default mode network (DMN), dorsal attention network (DAN), fronto-parietal network (FPN), and the salience network (SN) (Yeo et al., 2011). Synchronized activity within each network is observed under resting conditions and posited to underlie specific cognitive-affective functions including interception and attentional control (Hellyer et al., 2013). Converging evidence from other techniques suggests that neuromodulation can affect brain networks beyond the focal stimulation target (To et al., 2018; Pini et al., 2018; Santarnecchi et al., 2018). Thus, in order to fully elucidate the neuroplastic changes associated with tFUS, it is necessary to investigate ‘down-steam’ changes within and between well-characterized intrinsic neural networks.

In the present study, we collected resting-state BOLD fMRI before and after tFUS to characterize changes in rsFC. We used a sonication paradigm adapted from prior studies to inhibit/disrupt amygdala activity (Spivak et al., 2022; Folloni et al., 2019). We also included a second (within-subjects) target as an active regional comparison – the entorhinal cortex (ErC). The ErC is implicated in memory formation (Montchal et al., 2019) and hypoactive in diseases characterized by memory disturbance, e.g., Alzheimer’s disease and mild cognitive impairment (Igarashi, 2023). In a randomized, double-blind, within-subject crossover study design, we enrolled participants to complete two tFUS sessions, separated by a 14-day between-session window. The sonication paradigm for left ErC was hypothesized to excite/stimulate and therefore increase ErC activity and connectivity (Spivak et al., 2022; Dell’Italia et al., 2022). In a prior published study in these participants, we demonstrated that the sonication protocols selectively increased perfusion in the targeted region and not in the contralateral homolog or either bilateral control region (Kuhn et al., 2023). Data were collected and examined blindly to determine the target specific network changes in rsFC pre-to-post sonication. We broadly expected decreased rsFC connectivity within and between network nodes following sonication of the amygdala target and increased rsFC connectivity within and between network nodes following sonication of the ErC target based on the specific tFUS parameters.

## Methods

### Participants

A total of twenty healthy adults were enrolled in the study: 17 individuals completed both experimental sessions (4 scans), 18 individuals completed the amygdala tFUS session (2 scans), and 19 individuals completed the ErC tFUS session (2 scans). All participants were required to be between 30-85 years of age, right-handed, and proficient in English. Exclusionary criteria included contraindications for MRI, history of serious head injury, history of any major psychiatric illness requiring treatment, and history of any major neurological disorder (e.g., epilepsy) or serious illness (e.g., cancer). The age range was selected to obtain a sample representative of healthy adult aging to ultimately expand this line of work into this understudied population. In individuals 60 years and older, participants were required to score >30 on the Telephone Interview for Cognitive Status modified (TICS-M) (REF) to ensure absence of cognitive impairment.

The sample, on average, was 61.38 (7.75) years old, 56% female, and were 37% Caucasian American, 31% Latinx American, 19% African American and 13% Asian American. All procedures were in accordance with the Declaration of Helsinki and approved by the University of California, Los Angeles (UCLA) Institutional Review Board prior to enrollment. All participants provided written informed consent.

### Procedures

The study was a double-blind randomized, within-subjects crossover clinical trial (NCT03717922). Each participant completed two experimental sessions that were separated by exactly two weeks. Resting state FC was collected pre-tFUS and post-tFUS. Target order was randomized: one session targeted the right amygdala (experimental target) and the other session targeted the left entorhinal cortex (ErC; control target). Participants and study staff performing statistical analyses were blinded to the target (1 vs. 2). Following each lab session, participants were followed for three continuous days to access possible adverse events.

### MRI-Guided tFUS Sonication Protocol

The details of our sonication protocol are published elsewhere (Kuhn et al., 2023). In brief, sonications were delivered using a single-element transducer placed above the ear at the temporal window and targeted using real-time structural MRI navigation inside the scanner. The amygdala sonication protocol was designed to decrease or inhibit neural activity whereas the ErC sonication protocol was designed to increase or excite neural activity. Both paradigms used a 5% duty cycle, in 10 cycles of 30 seconds on, 30 seconds off, for a total of 5 minutes of non-consecutive tFUS. The amygdala target included a 5 ms pulse width repeated at a 10 Hz pulse repetition frequency (PRF) while the ErC target included a 0.5 ms pulse width repeated at a 100 Hz PRF. Across both sessions, the fundamental frequency was 0.65 MHz and the I_spta.3_ was 720 mW/cm^2^.

tFUS was performed inside the scanner. A 30-second SCOUT imaging sequence was used to visualize the transducer and its orthogonal line into the brain from the interface of the transducer and gel pad. The focal sonication depth was 65mm or 55mm (BrainSonix Corp., Sherman Oaks, CA, USA; Schafer et al., 2021) depending on each participants anatomical requirements to reach the desired brain target. The transducer was manually moved, as necessary, to correct its position for the appropriate target. The focus of the targeting line was either the centromedian aspect of the amygdala or the interface of the ErC and the perforant pathway.

### Resting State and Structural Data Acquisition

The MRI data were collected using a 3T Siemens MAGNETOM Prisma fit scanner (Siemens Medical Solution, Erlangen, Germany) located at the UCLA Center for Cognitive Neuroscience. A 20-channel head coil was used for all acquisition sequences to accommodate the tFUS transducer. Resting-state BOLD data were collected before and after tFUS using a GRE EPI sequence with the following acquisition parameters: TR = 800 ms, TE = 37 ms, flip angle = 52°, FOV = 208 mm (AP and RL) x 144 mm (FH), voxel size = 2.0 × 2.0 × 2.0 mm, slice thickness = 2.0 mm, slice count = 72 slices, phase encoding direction = AP, multi-band acceleration factor = 8, acquisition mode = interleaved, and total volumes = 488; thus, each resting-state run had a total run time of 390.40 seconds. Framewise Integrated Real-time MRI Monitoring (FIRMM) (Dosenbach et al., 2017) was used during the collection of all BOLD data to monitor for participant motion.

To correct for geometric distortions, opposite phase-encoded spin echo field map images were collected with the following parameters: phase encoding directions = AP and PA, TR = 8000 ms, TE = 66 ms, voxel size = 2.0 × 2.0 × 2.0 mm, FOV = 208 mm (AP and RL) and 144 mm (FH), slice thickness = 2.0 mm, flip angle = 90°, and refocus flip angle = 180°, and single-band acquisition. Additionally, structural MP-RAGE T1-weighted scans were acquired with the following parameters: orientation = sagittal, slices = 176, voxel size = 1.0 × 1.0 × 1.0 mm, slice thickness = 1.0 mm, TR = 2300 ms, TE = 2.98 ms, TI = 900 ms, flip angle = 9°, FOV = 256 mm (FH) and 248 mm (AP), and acceleration factor = 2 (GRAPPA).

### Data Preprocessing

All MRI data processing and analyses was carried out using the storage and computing service provided by the Ohio Supercomputer Center. The BOLD data were head motion corrected using mcflirt (FSL v6.0.4). Distortion correction was estimated from opposite phase-encoded spin-echo images using FSL’s TOPUP. After which, the BOLD data was co-registered to the subject’s T1w using boundary-based registration as implemented by bbregister (FreeSurfer v7.1.1) with 12 degrees of freedom. The BOLD data was then brought to the MNI template space by applying the T1w-to-MNI template (MNI152NLin2009cAsym) warp computed by antsRegistration (ANTs v2.3.5). To minimize smoothing effects, all the transformations were concatenated and applied in a single step to the BOLD volumes using antsApplyTransforms (ANTS v2.3.5); the images were sampled to the final space using Lanczos interpolation.

The preprocessed BOLD data were further denoised using the CONN toolbox (v21a). Effect of nuisance variables such as signals from white matter and cerebrospinal areas, movement parameters and their first derivative, and outlier scans were regressed out using the default implementation in CONN. A band-pass filter of 0.008 – 0.09 Hz, and linear detrending was also applied.

Nineteen regions of interest (ROI) were chosen from CONN’s network cortical ROIs atlas to be included in our analyses to estimate ROI-to-ROI functional connectivity. These regions included four ROIs from the default mode network, seven ROIs from the salience network, four ROIs from the frontal parietal network, and four ROIs from the dorsal attention network. The individual ROIs are listed in **Table 1**. A total of 171 ROI-to-ROI functional connections were estimated between the chosen regions. Here ROI-to-ROI functional connectivity was estimated as the Fisher transformed bivariate correlation coefficient between any given pair of ROI BOLD timeseries.

**Table 1.** Regions-of-interest used in the Primary Analyses.

| **Table 1.** Regions-of-interest used in the Primary Analyses |  |
| --- | --- |
| **Network** | **Region** |
| Dorsal Attention | IntraParietal Sulcus (IPS) (R) (39,-42,54) |
| Dorsal Attention | IntraParietal Sulcus (IPS) (L) (-39,-43,52) |
| Dorsal Attention | Frontal Eye Field (FEF) (R) (30,-6,64) |
| Dorsal Attention | Frontal Eye Field (FEF) (L) (-27,-9,64) |
| Salience | SupraMarginal Gyrus (SMG) (L) (-60,-39,31) |
| Salience | SupraMarginal Gyrus (SMG) (R) (62,-35,32) |
| Salience | Anterior Insula (AIC) (R) (47,14,0) |
| Salience | Anterior Insula (AIC) (L) (-44,13,1) |
| Salience | Anterior Cingulate (ACC) (0,22,35) |
| Salience | Rostral Prefrontal Cortex (RPFC) (L) (-32,45,27) |
| Salience | Rostral Prefrontal Cortex (RPFC) (R) (32,46,27) |
| Default Mode | Medial Prefrontal Cortex (MPFC) (1,55,-3) |
| Default Mode | Lateral Parietal (LP) (L) (-39,-77,33) |
| Default Mode | Lateral Parietal (LP) (R) (47,-67,29) |
| Default Mode | Precuneus Cortex (PCC) (1,-61,38) |
| Fronto Parietal | Posterior Parietal Cortex (PPC) (L) (-46,-58,49) |
| Fronto Parietal | Posterior Parietal Cortex (PPC) (R) (52,-52,45) |
| Fronto Parietal | Lateral Prefrontal Cortex (LPFC) (L) (-43,33,28) |
| Fronto Parietal | Lateral Prefrontal Cortex (LPFC) (R) (41,38,30) |

### Data Analysis Plan

We first performed a within-subjects paired samples *t*-test on all ROI-to-ROI connections for the experimental amygdala target and control ErC target, separately. Given that this was a preliminary investigation, to balance between statistical power and Type I and II error, we set a threshold of *p* < .01, uncorrected for multiple comparisons. Pre-to-post changes in rsFC that were found to be statistically significant (for either target) were then examined for target specificity. Extracted parameter estimates were entered into a target (amygdala vs. ErC) by time (pre vs. post) within-subjects repeated measures ANOVA. Significant two-way interactions were then followed-up by performing standard within-subjects comparisons.

## Results

### Amygdala Target: Network Changes Pre-to-Post Sonication

Results revealed significant change in ROI-to-ROI rsFC between several regions, across different networks. Results are displayed in Figure 1. The following pre-to-post sonication changes in connectivity were observed: 1) increase in left RPFC (SA network) to PCC (DMN network); 2) increase in right AIC (SA network) to PCC (DM network); 3) increase in right AIC (SA network) to left RPFC (SA network); and 4) decrease in right RPFC (SA network) to right LPFC (FP network).

**Figure 1.**
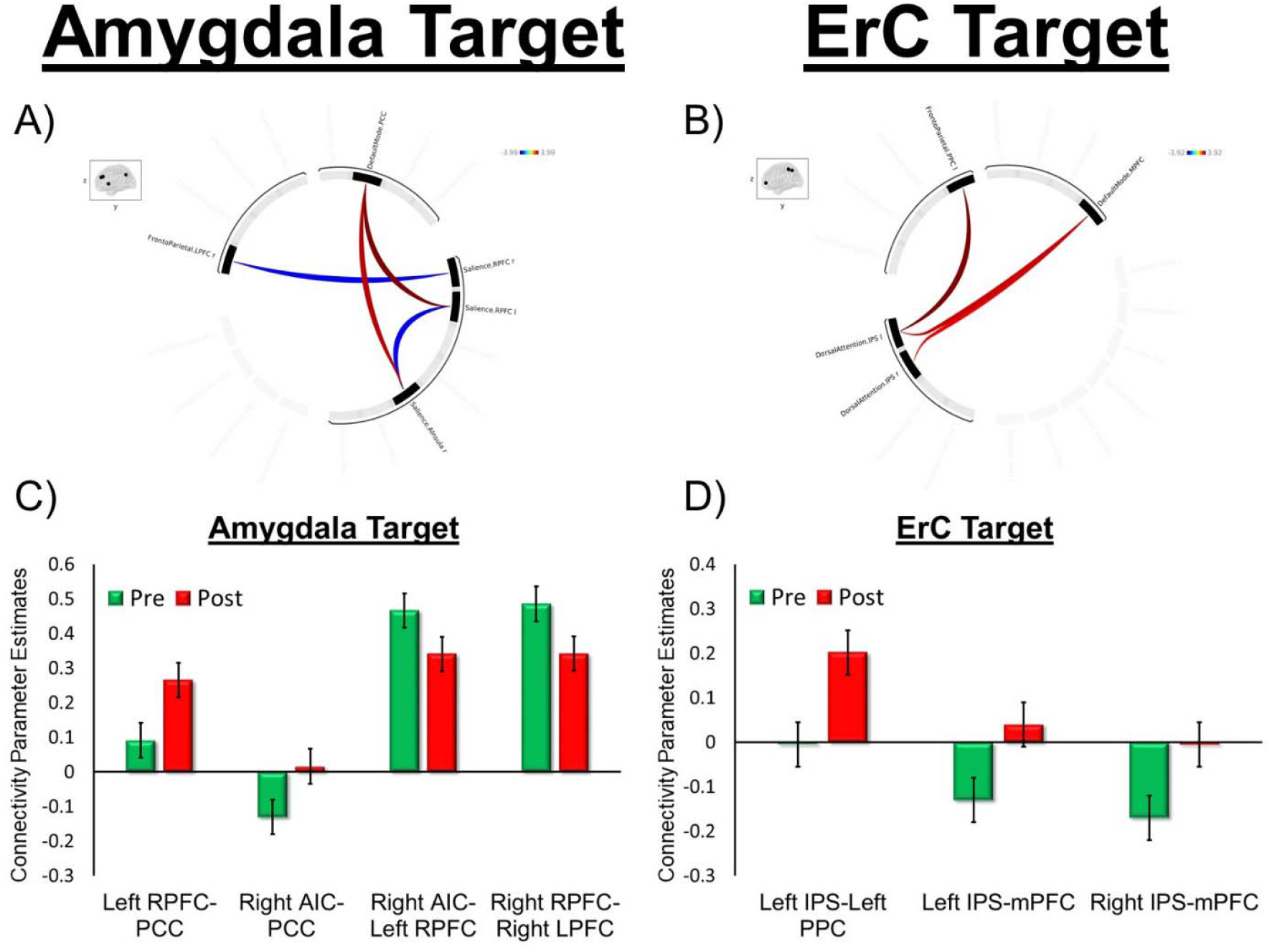
1A and 1B shows circular plots summarizing significant Region of Interest (ROI)-to-ROI connections in a paired test for the amygdala (Fig 1A) and entorhinal cortex (Fig 1B) targets, highlighting differences at p < 0.01. The plot was generated using the CONN toolbox. Significant connections are indicated by colored lines, with red lines representing connections that had greater connectivity post-tFUS compared to pre-tFUS and blue representing connections that had greater connectivity pre-tFUS compared to post-tFUS. The intensity of the colors correlates with the strength of the connections, as shown in the color bar. The inset shows the brain regions involved in the analysis. Fig 1A includes connections between the Default Mode Network (PCC), FrontoParietal Network (LPFC), and Salience Network (RPFC and Anterior Insula). Fig 1B includes connections between the Default Mode Network (mPFC), FrontoParietal Network (RPFC), and Dorsal Attention Network (IPS). Fig 1C and 1D display extracted connectivity parameter estimates from node connections that were found to significantly differ pre-to-post sonication.

### ErC Target: Network Changes Pre-to-Post Sonication

Results revealed significant increases in ROI-to-ROI rsFC between the DA network and the FP and DM networks. The following pre-to-post sonication changes in connectivity were observed: 1) increase in left IPS (DA network) to left PPC (FP network); 2) increase in left IPS (DA network) and mPFC (DM network); and 3) increase in right IPS (DA network) to mPFC (DM network).

### Target Comparison

rsFC parameter estimates for the seven ROI-to-ROI connections (identified above; 4 amygdala target findings and 3 ErC target findings) were extracted for both targets, pre- and post-sonication for the 17 individuals that completed all 4 scans. These parameter estimates were then entered into a target (amygdala vs. ErC) by time (pre-vs. post-stimulation) repeated measures ANOVA to assess the specificity of each finding.

The results of the seven ANOVAs are presented in **Table 2**. There was a significant target by time interaction on right RPFC (SA network) to right LPFC (FP network) rsFC. Sonication of the right amygdala resulted in a decrease in right RPFC-right LPFC rsFC (*t* [16] = 2.71, *p* < 0.01); however, sonication of the left ErC produced no change in connectivity between these ROIs (*t* [16] = -1.56, *p* = 0.14). There was also a target by time interaction on left IPS (DA network) to mPFC (DM network) and right IPS (DA network) to mPFC (DM network) rsFC. Sonication of the left ErC resulted in increases in left IPS to mPFC (*t* [16] = -3.14, *p* < 0.01) and right IPS to mPFC (*t* [16] = -2.99, *p* = 0.01). There were no changes in left IPS to mPFC (*t* [16] = 0.72, *p* = 0.48) nor right IPS to mPFC (*t* [16] = 0.77, *p* = 0.45) following sonication of the right amygdala.

**Table 2.**
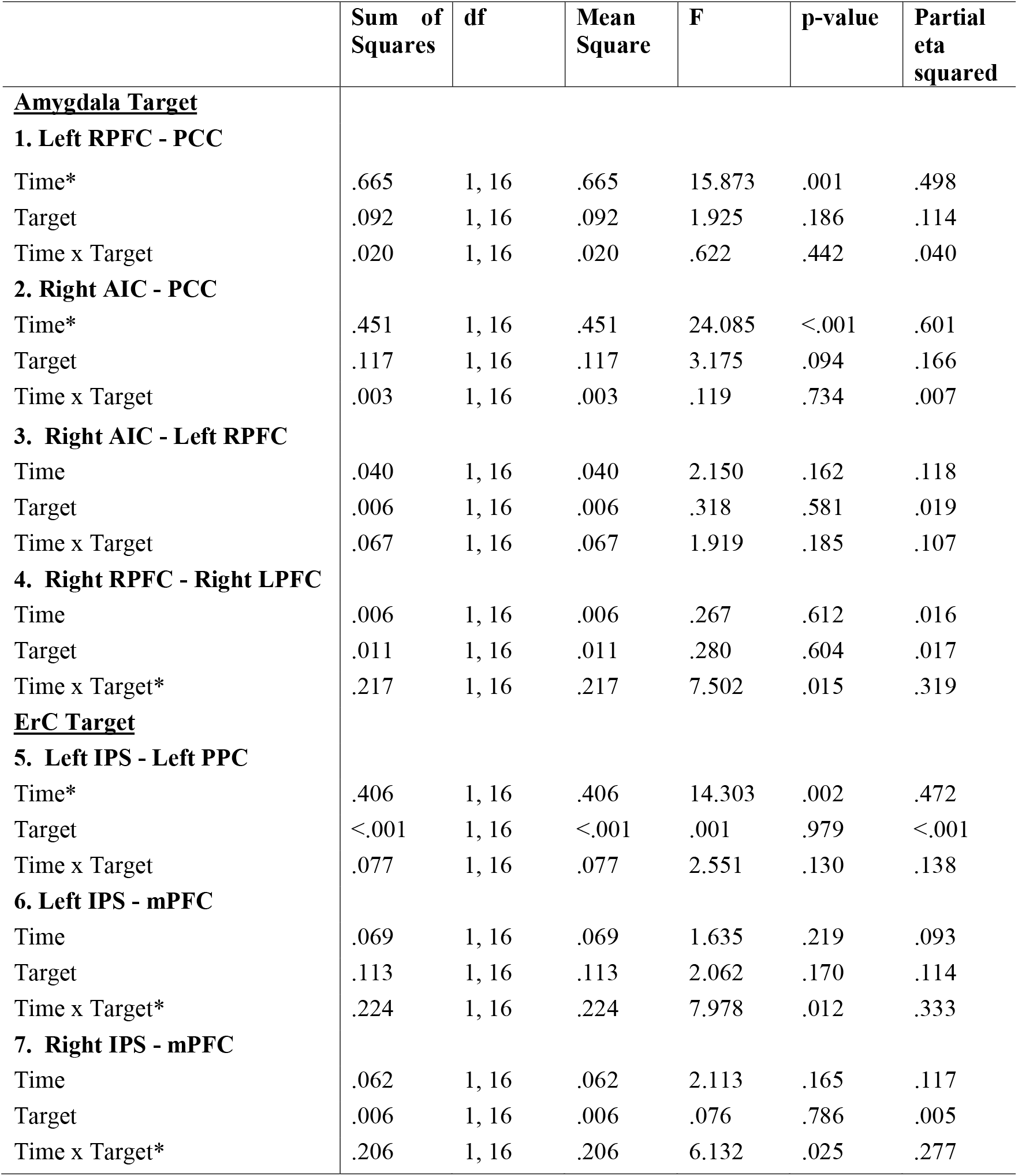
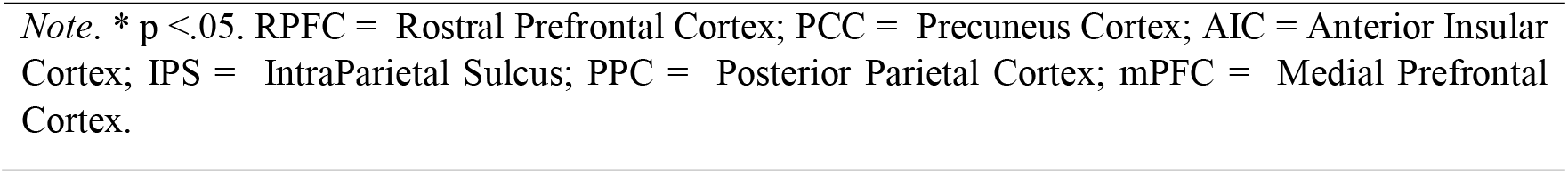
Results of the repeated measures analysis of variance for each sonication target.

## Discussion

The current study used resting-state BOLD fMRI data before and after tFUS to examine changes in functional connectivity between nodes of large-scale rs-networks. The results revealed that tFUS of the amygdala (and ErC) changed connectivity within and between rs-networks. Pre-to-post sonication of the right amygdala was found to modulate connectivity within nodes of the SAN and between nodes of the SAN and the DMN and FPN. A decrease in SAN to FPN connectivity was specific to the amygdala target. Pre-to-post sonication of the left ErC was found to modulate connectivity between the DAN and FPN and DMN. An increase in DAN to DMN connectivity was specific to the ErC target. These preliminary findings suggest that tFUS may result in neuroplastic changes outside of the focal neural target.

tFUS of the amygdala was associated with changes in connectivity within the SAN and between the SAN and the DMN and FPN. The SAN is involved in the integration of emotionally salient information and makes inferences regarding interoceptive awareness, threat/reward value, and outcome probabilities to appropriately guide approach and avoidance behavior (Seeley et al. 2007; Vytal & Hamann 2010). Related, the functional integrity within the SAN underlies the subjective experience of affective states (De Witte et al., 2017; Prillwitz et al., 2018). Theory and research further show that the SAN interacts with other large-scale rs-networks and plays a critical role in the dynamic switching between the DMN and the FPN (Sridharan et al., 2008; Goulden et al., 2014). This switching function allows for efficient engagement and disengagement of goal-directed resources. Upon detection of a salient event, the SAN facilities access to attention and working memory via the FPN (Cai et al., 2021). During rest, the SAN engages the DMN, which supports basic self-referential functions and internally focused attention (Yoon et al., 2019). Regarding the present findings, acute tFUS-related increases in rsFC within the SAN may reflect increased salience processing. Meanwhile, increased SAN to DMN rsFC and decreased SAN to FPN rsFC may signal changes in the SAN switching functions between networks. The decreased rsFC connectivity between the SAN and FPN was unique to the amygdala target and thus, amygdala inhibition may uniquely decrease SAN input to the FPN. Numerous studies have demonstrated that anxiety and other stress-related disorders are characterized by aberrant SAN function including deficient network switching (e.g., Li et al. 2023). Additional studies are therefore critically needed to replicate these preliminary findings and elucidate how tFUS-related SAN changes may impact anxiety symptoms.

tFUS of the ErC was used as an active within-subjects comparison. Sonication of the ErC resulted in pre-to-post changes in DAN to FPN and DMN rsFC. The DAN is comprised of regions in the frontal and parietal cortices (Osher, Brissenden, & Somers, 2019), which become engaged when attention is voluntarily shifted to salient objects and/or locations (Corbetta & Shulman, 2009) and during visual exploration (Corbetta et al., 1998). The DAN also underlies attentional control via top-down influences on the visual cortex (Osher, Brissenden, & Somers, 2019. Anti-correlation between the DAN and DMN is characteristic of typical brain function (Fox et al., 2009, Kelly et al., 2008, Kundu et al., 2013) which reflects the distinct attentional processes these two networks serve: the DMN mediates internally directed attention whereas the DAN mediates externally directed attention (Spreng, Shoemaker, & Turner, 2017). The magnitude of anti-correlation between the DAN and DMN has been associated with performance of attention-based tasks in healthy young adults (Hampson et al., 2010); though this association is attenuated by age and seems to differ in patients (Wang et al., 2019; Spreng et al., 2016; Owens et al., 2020). In at least one prior study in adults with depression, positive connectivity between the DAN and DMN was associated with increased memory accuracy for objects (Satz et al., 2022). Interestingly, in a sample of older adults, we found that sonication of the ErC resulted in increases in DAN to DMN rsFC, and this increase was specific to the ErC target. We also found increases in DAN to FPN rsFC which are two networks that functionally interact to support perceptual attention (Dixon et al., 2018). Although the functional significance of these acute changes are unknown, it is noteworthy that the DAN, DMN, and FPN are all involved in the cognitive processes that mediate learning and memory – core functions of the ErC target. Thus, the ErC as a target for tFUS in patient groups with amnestic syndromes appears to warrant further exploration.

The pattern of results indicates that some FC changes were more robust following sonication of one active target but not the other. Sonication of the amygdala target using an inhibition protocol resulted in decreases in rsFC between the SAN and FPN. Meanwhile, sonication of the ErC target using an excitation protocol resulted in increases in DAN to DMN rsFC. These findings are broadly consistent with our study hypotheses and demonstrate that tFUS can have ‘downstream’ effects that alter rsFC patterns within and between large-scale networks. Other neuromodulation techniques such as deep brain stimulation and transcranial magnetic stimulation have shown similar extended effects (Haslinger et al., 2003; Coenen et al., 2014; Esser, Hill, & Tononi, 2005). It is noteworthy that changes involving the SAN were exclusively found with tFUS of the amygdala, given that the SAN is the core rs-network involved in salience and emotion processing, and is implicated in the pathophysiology of anxiety disorders (Pannekoeket al., 2013). Investigating if and how these SAN-level changes influence affective states and anxiety symptoms is a critical next step, particularly given existing research highlighting the potential clinical utility of tFUS of the amygdala for anxiety (Chou et al., 2024). Meanwhile, the ErC is a region of the brain involved in memory (Guzman et al., 2013; Eichenbaum, 2000) and tFUS of the ErC modulated DAN rsFC with other attention networks. Attention and memory disturbance, and associated hypoactivity of the ErC, underliesseveral neurological disorders, including Alzheimer’s disease (Donix et al., 2013, Maass et al., 2018). Given the dissociative findings observed here, investigation into the functional significance of DAN-level changes as they pertain to cognitive processes like memory may also be a fruitful next step.

The current study had many strengths including the double-blind, within-subject crossover design. The study also had several important limitations. First, the study focused on healthy older adults given that there is a need for novel psychiatric and neurological therapeutic strategies for this developmental period. Prior research shows that there are age-related changes in rsFC between large-scale networks including those investigated in the current study (Grady et al., 2016; Meunier et al., 2009). It is therefore unclear whether the present findings would generalize to younger adults. Second, the sample size was small; although consistent with other published studies involving tFUS in humans (Kuhn et al., 2023; Pasquinelli et al., 2019). It is possible the study was under-powered to detect certain target-specific network level changes. Related, a total of 171 ROI-to-ROI connections were initially tested without stringent correction for multiple comparisons. The current findings are therefore considered preliminary and require replication. Lastly, the study focused on immediate, acute changes (pre-to-post sonication) and did not measure real-time functional outcomes. Additional studies are needed to determine the duration and durability of the observed findings and if and how network-level changes relate to cognitive and emotional outcomes.

The present findings add to a growing body of literature on the feasibility of tFUS and its acute neural effects. Converging research, including our own, show that tFUS can target desired areas of the deep brain without engaging nearby structures (Kuhn et al., 2023). We also demonstrate that tFUS can evoke broader changes, beyond the focal sonication target. There are many important next steps in this line of work including investigation of functional significance and therapeutic value of these network changes.

## References

Cai, W., Ryali, S., Pasumarthy, R., Talasila, V., & Menon, V. (2021). Dynamic causal brain circuits during working memory and their functional controllability. Nature communications, 12(1), 3314. 10.1038/s41467-021-23509-x

Coenen, V. A., Allert, N., Paus, S., Kronenbürger, M., Urbach, H., & Mädler, B. (2014). Modulation of the cerebello-thalamo-cortical network in thalamic deep brain stimulation for tremor: a diffusion tensor imaging study. Neurosurgery, 75(6), 657–670. 10.1227/NEU.0000000000000540

Chou, T., Deckersbach, T., Guerin, B., Sretavan Wong, K., Borron, B. M., Kanabar, A., Hayden, A. N., Long, M. P., Daneshzand, M., Pace-Schott, E. F., & Dougherty, D. D. (2024). Transcranial focused ultrasound of the amygdala modulates fear network activation and connectivity. Brain stimulation, 17(2), 312–320. Advance online publication. 10.1016/j.brs.2024.03.004

De Witte, N.A.J., Mueller, S.C. White matter integrity in brain networks relevant to anxiety and depression: evidence from the human connectome project dataset. Brain Imaging and Behavior 11, 1604–1615 (2017). 10.1007/s11682-016-9642-2

Dell’Italia, J., Sanguinetti, J. L., Monti, M. M., Bystritsky, A., & Reggente, N. (2022). Current state of potential mechanisms supporting low intensity focused ultrasound for neuromodulation. Frontiers in human neuroscience, 16, 872639. 10.3389/fnhum.2022.872639

di Biase, L., Falato, E., Caminiti, M. L., Pecoraro, P. M., Narducci, F., & Di Lazzaro, V. (2021). Focused Ultrasound (FUS) for Chronic Pain Management: Approved and Potential Applications. Neurology research international, 2021, 8438498. 10.1155/2021/8438498

Donix, M., Burggren, A. C., Scharf, M., Marschner, K., Suthana, N. A., Siddarth, P., Krupa, A. K., Jones, M., Martin-Harris, L., Ercoli, L. M., Miller, K. J., Werner, A., von Kummer, R., Sauer, C., Small, G. W., Holthoff, V. A., & Bookheimer, S. Y. (2013). APOE associated hemispheric asymmetry of entorhinal cortical thickness in aging and Alzheimer’s disease. Psychiatry research, 214(3), 212–220. 10.1016/j.pscychresns.2013.09.006

Eichenbaum, H. (2000). A cortical–hippocampal system for declarative memory. Nature reviews neuroscience, 1(1), 41–50.

Esser, S. K., Hill, S. L., & Tononi, G. (2005). Modeling the effects of transcranial magnetic stimulation on cortical circuits. Journal of neurophysiology, 94(1), 622–639. 10.1152/jn.01230.2004

Folloni, D., Verhagen, L., Mars, R. B., Fouragnan, E., Constans, C., Aubry, J. F., Rushworth, M.F. S., & Sallet, J. (2019). Manipulation of Subcortical and Deep Cortical Activity in the Primate Brain Using Transcranial Focused Ultrasound Stimulation. Neuron, 101(6), 1109–1116.e5. 10.1016/j.neuron.2019.01.019

Gerin, M. I., Viding, E., Pingault, J. B., Puetz, V. B., Knodt, A. R., Radtke, S. R., Brigidi, B. D., Swartz, J. R., Hariri, A. R., & McCrory, E. J. (2019). Heightened amygdala reactivity and increased stress generation predict internalizing symptoms in adults following childhood maltreatment. Journal of child psychology and psychiatry, and allied disciplines, 60(7), 752–761. 10.1111/jcpp.13041

Guzman, V. A., Carmichael, O. T., Schwarz, C., Tosto, G., Zimmerman, M. E., Brickman, A. M., & Alzheimer’s Disease Neuroimaging Initiative. (2013). White matter hyperintensities and amyloid are independently associated with entorhinal cortex volume among individuals with mild cognitive impairment. Alzheimer’s & Dementia, 9(5), S124–S131.

Haslinger, B., Boecker, H., Büchel, C., Vesper, J., Tronnier, V. M., Pfister, R., Alesch, F., Moringlane, J. R., Krauss, J. K., Conrad, B., Schwaiger, M., & Ceballos-Baumann, A. O. (2003). Differential modulation of subcortical target and cortex during deep brain stimulation. NeuroImage, 18(2), 517–524. 10.1016/s1053-8119(02)00043-5

Hellyer, P. J., Shanahan, M., Scott, G., Wise, R. J., Sharp, D. J., & Leech, R. (2014). The control of global brain dynamics: opposing actions of frontoparietal control and default mode networks on attention. The Journal of neuroscience : the official journal of the Society for Neuroscience, 34(2), 451–461. 10.1523/JNEUROSCI.1853-13.2014

Igarashi K. M. (2023). Entorhinal cortex dysfunction in Alzheimer’s disease. Trends in neurosciences, 46(2), 124–136. 10.1016/j.tins.2022.11.006

Kuhn, T., Spivak, N. M., Dang, B. H., Becerra, S., Halavi, S. E., Rotstein, N., Rosenberg, B. M., Hiller, S., Swenson, A., Cvijanovic, L., Dang, N., Sun, M., Kronemyer, D., Berlow, R., Revett, M. R., Suthana, N., Monti, M. M., & Bookheimer, S. (2023). Transcranial focused ultrasound selectively increases perfusion and modulates functional connectivity of deep brain regions in humans. Frontiers in neural circuits, 17, 1120410. 10.3389/fncir.2023.1120410

Li, R., Shen, F., Sun, X., Zou, T., Li, L., Wang, X., … & Chen, H. (2023). Dissociable salience and default mode network modulation in generalized anxiety disorder: a connectome-wide association study. Cerebral Cortex, 33(10), 6354–6365.

Moosa, S., Martínez-Fernández, R., Elias, W. J., Del Alamo, M., Eisenberg, H. M., & Fishman, P. S. (2019). The role of high-intensity focused ultrasound as a symptomatic treatment for Parkinson’s disease. Movement disorders : official journal of the Movement Disorder Society, 34(9), 1243–1251. 10.1002/mds.27779

Maass, A., Lockhart, S. N., Harrison, T. M., Bell, R. K., Mellinger, T., Swinnerton, K., Baker, S. L., Rabinovici, G. D., & Jagust, W. J. (2018). Entorhinal Tau Pathology, Episodic Memory Decline, and Neurodegeneration in Aging. The Journal of neuroscience : the official journal of the Society for Neuroscience, 38(3), 530–543. 10.1523/JNEUROSCI.2028-17.2017

Ohman A. (2005). The role of the amygdala in human fear: automatic detection of threat. Psychoneuroendocrinology, 30(10), 953–958. 10.1016/j.psyneuen.2005.03.019

Osher, D. E., Brissenden, J. A., & Somers, D. C. (2019). Predicting an individual’s dorsal attention network activity from functional connectivity fingerprints. Journal of neurophysiology, 122(1), 232–240. 10.1152/jn.00174.2019

Owens, M. M., Yuan, D., Hahn, S., Albaugh, M., Allgaier, N., Chaarani, B., Potter, A., & Garavan, H. (2020). Investigation of Psychiatric and Neuropsychological Correlates of Default Mode Network and Dorsal Attention Network Anticorrelation in Children. Cerebral cortex (New York, N.Y. : 1991), 30(12), 6083–6096. 10.1093/cercor/bhaa143

Pannekoek, J. N., Veer, I. M., van Tol, M. J., van der Werff, S. J., Demenescu, L. R., Aleman, A., Veltman, D. J., Zitman, F. G., Rombouts, S. A., & van der Wee, N. J. (2013). Resting-state functional connectivity abnormalities in limbic and salience networks in social anxiety disorder without comorbidity. European neuropsychopharmacology : the journal of the European College of Neuropsychopharmacology, 23(3), 186–195. 10.1016/j.euroneuro.2012.04.018

Pasquinelli C., Hanson L. G., Siebner H. R., Lee H. J., Thielscher A. (2019). Safety of transcranial focused ultrasound stimulation: A systematic review of the state of knowledge from both human and animal studies. Brain Stimul. 12 1367–1380.

Pini, L., Manenti, R., Cotelli, M., Pizzini, F. B., Frisoni, G. B., & Pievani, M. (2018). Non-Invasive Brain Stimulation in Dementia: A Complex Network Story. Neuro-degenerative diseases, 18(5-6), 281–301. 10.1159/000495945

Prillwitz, C. C., Rüber, T., Reuter, M., Montag, C., Weber, B., Elger, C. E., & Markett, S. (2018). The salience network and human personality: Integrity of white matter tracts within anterior and posterior salience network relates to the self-directedness character trait. Brain research, 1692, 66–73. 10.1016/j.brainres.2018.04.035

Ressler, K. J., & Nemeroff, C. B. (2000). Role of serotonergic and noradrenergic systems in the pathophysiology of depression and anxiety disorders. Depression and anxiety, 12 Suppl 1, 2–19. 10.1002/1520-6394(2000)12:1+<2::AID-DA2>3.0.CO;2-4

Romanella, S. M., Sprugnoli, G., Ruffini, G., Seyedmadani, K., Rossi, S., & Santarnecchi, E. (2020). Noninvasive Brain Stimulation & Space Exploration: Opportunities and Challenges. Neuroscience and biobehavioral reviews, 119, 294–319. 10.1016/j.neubiorev.2020.09.005

Santarnecchi, E., Momi, D., Sprugnoli, G., Neri, F., Pascual-Leone, A., Rossi, A., & Rossi, S. (2018). Modulation of network-to-network connectivity via spike-timing-dependent noninvasive brain stimulation. Human brain mapping, 39(12), 4870–4883. 10.1002/hbm.24329

Stern J. M., Spivak N. M., Becerra S. A., Kuhn T. P., Korb A. S., Kronemyer D., et al. (2021). Safety of focused ultrasound neuromodulation in humans with temporal lobe epilepsy. Brain Stimul. 14 1022–1031.

Spivak N. M., Sanguinetti J. L., Monti M. M. (2022). Focusing in on the future of focused ultrasound as a translational tool. Brain Sci. 12:158.

Spreng, R. N., Shoemaker, L., & Turner, G. R. (2017). Executive functions and neurocognitive aging. In E. Goldberg (Ed.), Executive functions in health and disease (pp. 169–196). Elsevier Academic Press. 10.1016/B978-0-12-803676-1.00008-8

Spreng, R. N., Stevens, W. D., Viviano, J. D., & Schacter, D. L. (2016). Attenuated anticorrelation between the default and dorsal attention networks with aging: evidence from task and rest. Neurobiology of aging, 45, 149–160. 10.1016/j.neurobiolaging.2016.05.020

Tyler, W. J., Tufail, Y., Finsterwald, M., Tauchmann, M. L., Olson, E. J., & Majestic, C. (2008). Remote excitation of neuronal circuits using low-intensity, low-frequency ultrasound. PloS one, 3(10), e3511. 10.1371/journal.pone.0003511

Yeo BT, Krienen FM, Sepulcre J, Sabuncu MR, Lashkari D, Hollinshead M, Roffman JL, Smoller JW, Zollei L, Polimeni JR, Fischl B, Liu H, Buckner RL. The organization of the human cerebral cortex estimated by intrinsic functional connectivity. J Neurophysiol. 2011;106(3):1125–65.

To, W. T., De Ridder, D., Hart Jr, J., & Vanneste, S. (2018). Changing brain networks through non-invasive neuromodulation. Frontiers in human neuroscience, 12, 128.

Corbetta, M., and Shulman, G. L. (2002). Control of goal-directed and stimulus-driven attention in the brain. Nat. Rev. Neurosci. 3, 201–215.

Wang, J., Liu, J., Wang, Z., Sun, P., Li, K., & Liang, P. (2019). Dysfunctional interactions between the default mode network and the dorsal attention network in subtypes of amnestic mild cognitive impairment. Aging, 11(20), 9147–9166. 10.18632/aging.102380

Yoon, H. J., Seo, E. H., Kim, J. J., & Choo, I. H. (2019). Neural Correlates of Self-referential Processing and Their Clinical Implications in Social Anxiety Disorder. Clinical psychopharmacology and neuroscience : the official scientific journal of the Korean College of Neuropsychopharmacology, 17(1), 12–24. 10.9758/cpn.2019.17.1.12

